# Transarterial Embolization Using a Liquid Embolic Enhances Tumor Necrosis to Enable Improved Efficacy Compared with a Particle Embolic in a Translational Rat Model of Hepatocellular Carcinoma

**DOI:** 10.1101/2024.05.23.595637

**Authors:** Alexey Gurevich, Ariful Islam, Jonathan Wakim, Eva Yarsky, Ryan Kiefer, Ryan El-Ghazal, George McClung, David Peter Cormode, Gregory Jon Nadolski, Rony Avritscher, Stephen James Hunt, Terence Peter Ferrante Gade

## Abstract

**Purpose:** To compare the effectiveness of transarterial embolization (TAE) using a liquid embolic (LE) to TAE using particle embolics (PE) with respect to tumor ischemia and treatment response in a translational rat model.

**Materials and Methods:** HCC was induced in male Wistar rats using diethylnitrosamine. Rats were randomized into three treatment groups: sham transarterial embolization (TAE), TAE with LE, and TAE with PE. Tumor response to TAE was monitored via serial T2-weighted MRI scans and assessed using RECIST. Tumor necrosis and hypoxia were assessed through hematoxylin and eosin staining and pimonidazole immunohistochemistry, respectively. Statistical analyses were performed using chi-square tests, Kaplan-Meier estimates, and one-way ANOVAs, with significance set at p<0.05.

**Results:** Twenty-nine rats were randomized to TAE with LE (n=13), PE (n=13), or sham (n=3). TAE with LE demonstrated a significantly higher objective response rate (83%) compared to TAE with PE (28%; χ2 = 11.25, P=0.0008). Complete responses were observed in 50% of the LE-treated tumors versus 10% in the PE-treated group. LE treatment prolonged local progression-free survival (hazard ratio = 0.23; p=0.0085). Histological analysis, with an additional 14 rats randomized to TAE with LE (n=5), PE (n=6), or sham (n=3), showed greater necrosis and hypoxia in LE-treated tumors compared to PE. LE induced a significant reduction in viable tumor tissue (p<0.05) and increased necrotic and hypoxic tissue areas compared to PE (p<0.01).

**Conclusion:** LE significantly enhanced the therapeutic efficacy of TAE in a rat model of HCC compared to conventional PE. The results highlight LE’s potential to improve ischemia and necrosis, thereby offering a promising option improving the efficacy of embolization for HCC treatment.

## Introduction

Hepatocellular carcinoma (HCC) is the fifth leading cause of cancer death in the United States, and transarterial chemoembolization (TACE) continues to play a central role in the treatment for intermediate unresectable HCC. However, outcomes have been limited by variable response rates and disease recurrence (1–3). Suboptimal outcomes have been demonstrated to issue, at least in part, to incomplete embolization resulting from particle-microvasculature size mismatch and poor particle embolic (PE) penetration into the tumor vascular bed. These limitations lead to heterogeneous induction of ischemia and revascularization of the tumor feeding vessels (4–6).

These challenges have motivated the development of novel embolic agents to optimize vascular penetration and improve the degree of ischemia while limiting revascularization. Among these, liquid embolics (LE) have received considerable interest based on their potential to achieve capillary-level penetration and to enable complete occlusion by fully obstructing the target vasculature(7). LE formulations developed to date, including cyanoacrylates and ethylene-vinyl alcohol copolymers, have been limited by an inability to control polymerization rendering them susceptible to nontarget embolization, adherence of the LE to the catheter tip, and/or catheter blockage while also limiting the penetration to intratumoral arterioles and capillaries(8–10). The development of novel LE formulations may overcome this limitation by enabling delayed polymerization. The purpose of this study was to compare the efficacy of a next generation LE to a PE in a translational rat model of hepatocellular carcinoma (11).

## Materials and Methods

### Autochthonous Rat HCC Model of Transarterial Embolization

Animal studies were conducted in accordance with institutionally approved IRB protocols by adhering to Institutional Animal Care and Use Committee (IACUC) guidelines. As described previously, HCCs were induced in male Wistar rats weighing 300–400 g (Charles River Laboratories) through ad libitum oral administration of 0.01% diethylnitrosamine for 12 weeks, which induces autochthonous liver tumors through sequential progression of hepatitis, cirrhosis, and carcinogenesis (12). Rats were monitored for the development of HCCs using serial T2-weighted MRI scans (Agilent 4.7-T 40-cm horizontal bore MRI; repetition time, 1.4 seconds [respiratory-gated]; echo time, 59.1 msec) Rats bearing tumors measuring 100-200 mm in volume were randomized for experiment. Transarterial embolization (TAE) was performed using a transfemoral approach (13). Fluoroscopy and digital subtraction arteriography for TAE was performed using Cios Alpha angiographic imaging system (Siemens). Rats were euthanized when the TAE-targeted tumor, or another tumor reached established size criteria.

A custom-designed coaxial catheter system enabled successful selective delivery of liquid embolic into tumor feeding vessels (Figure 1A-E). TAE with LE was performed using the Embrace Hydrogel Embolic System (HES; Instylla LLC, Bedford, MA) which consists of two low viscosity liquid precursors, a polymer and an initiator, which, when combined in the target vessel, form a resorbable, water-based hydrogel. Precursors were initially mixed with contrast media for radiopacity as per the manufacturer’s instructions. LE was delivered under fluoroscopic guidance via this coaxial microcatheter system using a PHD Ultra Syringe Pump (Harvard Apparatus, Holliston, MA) at a rate of 75 µL/min., with the outer 1.7F microcatheter carrying the low viscosity initiator precursor molecules, and the inner 0.9F microcatheter carrying the low viscosity polymer precursor molecules. This coaxial, pump-controlled delivery system prevented premature precursor mixing and catheter plugging and non-target embolization. TAE using a PE was performed through fluoroscopically-guided injection of 40 µm Embozene (Varian, Palo Alto, CA) delivered through a 1.7F microcatheter using a Syringe Pump at 50 to 100 µL/min as described previously(13,14). Embolization using both LE and PE was performed to stasis.

**Figure 1:**
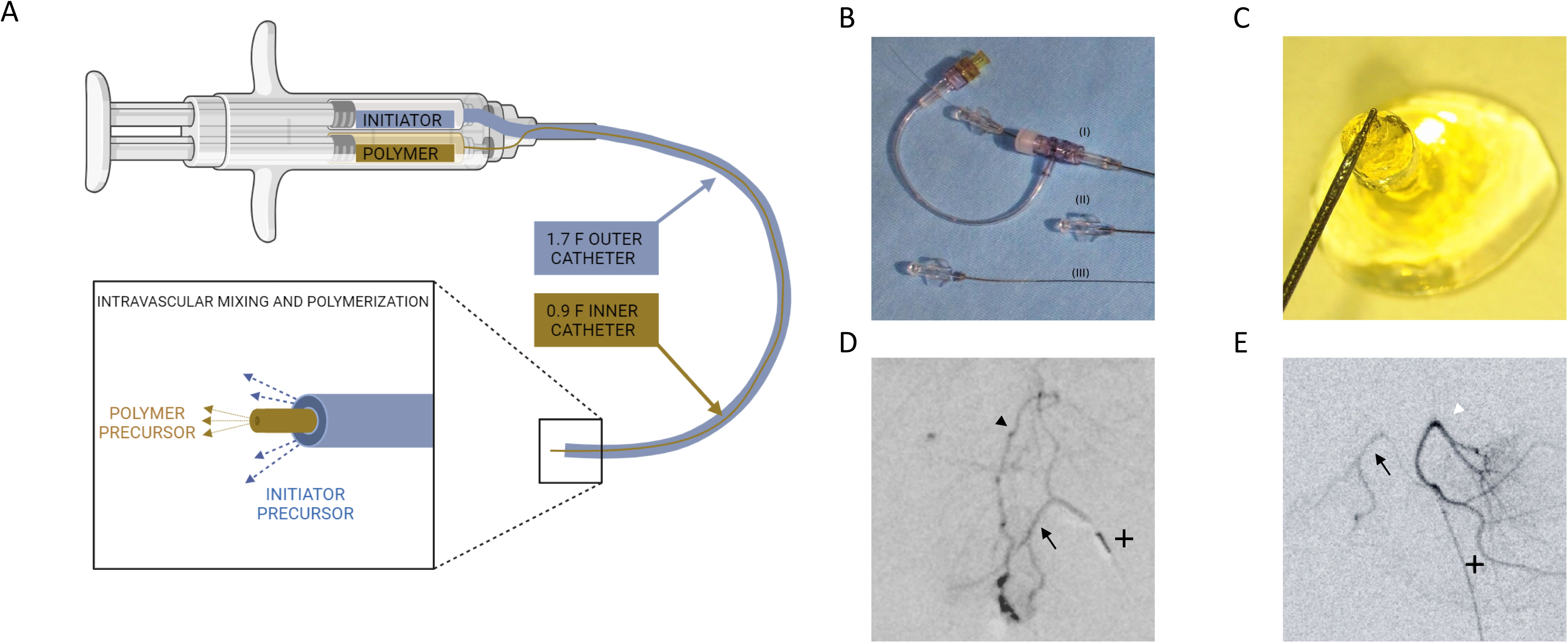
Dual lumen coaxial system used to deploy the hydrogel embolic. . (a) Dual catheter injection system allowed for simultaneous delivery of the polymer and initiator precursors, preventing mixing prior to entering the target vessel. (b) Photographs of the assembled coaxial system (I) along with the outer 1.7F catheter (II) and the inner 0.9F catheter (III) used to separate the two precursors for simultaneous delivery. (c) Gel polymerization upon mixing of the precursors as they exit the catheters. (d) Digital subtraction angiography prior to embolization demonstrating the catheter system within the common hepatic artery (cross) with perfusion to the right hepatic artery (black arrow) and the tumor feeding left hepatic artery (black arrowhead). (e) Digital subtraction angiography after embolization demonstrating the catheter system within the celiac artery (cross) with perfusion of the right hepatic artery (black arrow) and the left gastric artery (white arrowhead) as well as absence of perfusion to the left hepatic artery and tumor.

### MR Image Analysis

To assess efficacy, 29 male Wistar rats bearing HCCs measuring 100 to 200 mm were randomized to three groups including (i) sham TAE, (ii) TAE with LE and (iii) TAE with PE. Target tumor volume(s) for each rat were measured using ITK-snap segmentation software and averaged for each imaging session(15). Thirteen rats with 18 targetable tumors on MRI were treated with TAE using the LE, thirteen rats with 18 targetable tumors on MRI were treated with TAE using a PE, and 3 rats with 6 targetable tumors on MRI were treated with sham saline therapy. Responses were determined using these measurements and RECIST (16). Gompertzian transformed tumor growth kinetics were analyzed using linear regressions of MRI volumes (17). A separate cohort of 14 additional rats underwent TAE with liquid embolic (5 rats with 8 tumors), PE (6 rats with 10 tumors), or sham saline (3 rats with 5 tumors) embolization followed by IV administration of pimonidazole approximately 72 hours later. These rats were then euthanized, and target tumor tissues were processed for histological analyses.

### Histology

Treated and untreated tumors were harvested from rats following euthanasia in accordance with institutionally approved protocols Tumors were hemisected, fixed overnight in formalin, and dehydrated with 70% alcohol before paraffinization. The pathologic diagnosis of HCC was confirmed on 4-µm-thick hematoxylin and eosin–stained (H&E) sections by a hepatobiliary pathologist with 35 years of experience. Standard IHC protocols were used to stain tissues fixed in formalin and embedded in paraffin. Immunohistochemical staining was performed for alpha smooth muscle actin (a-SMA, Abcam, ab5694) and pimonidazole (Hypoxyprobe) as per the manufacturer’s instructions. Lugol’s Iodine staining was performed as described previously (18). Automated image analysis was performed on pimonidazole stained slides using ImageJ version 1.54h in a blinded fashion, wherein digitally scanned slides were anonymized with use of a unique identifier(19). Staining was quantified using Huang analysis to calculate the total percent tumor hypoxia and total image size(20). An overlay of the viable tissue identified on H&E staining was created and similarly quantified. Using these metrics, the total percent viable tumor, percent necrotic tumor, and percent hypoxic tumor were calculated.

### Statistical Analysis

Graphs and analyses were generated and performed using GraphPad Prism 9.4.1 (GraphPad Software). Statistical tests used a p-value <.05 to determine statistical significance. Normally distributed data are reported as the mean ± SD.

## Results

### TAE with Liquid Embolic Demonstrates Improved Therapeutic Efficacy

Rats bearing tumors were randomized into (i) sham, (ii) PE TAE and (iii) LE TAE groups with average pre-treatment volumes measuring 150.9 ±66.1 mm^3^ , 135.1 ±72.38 mm 150.5 ±98.1 mm on MRI, respectively. Volumetric assessments of post-TAE responses on T2-weighted MRI revealed improved efficacy for LE, as compared to PE treated tumors (Figure 2A-C). LE treated tumors demonstrated complete responses (CR) in 50% of tumors (9/18), partial responses (PR) in 33% of tumors (6/18) and progressive disease (PD) in 17% of tumors (3/18, Figure 2D). The PE treated group demonstrated CR in 11% of tumors (2/18), PR in 17% of tumors (3/18), stable disease (SD) in 39% of the tumors (7/18), and PD in 33% of the tumors (6/18, Figure 2E). Consistent with these findings, LE treated tumors demonstrated a significantly higher objective response rate of 83% (15/18) as compared to 28% for PE treated tumors (5/18; χ2 = 11.25, P=0.0008; Figure 2F). Kaplan-Meier estimates of local progression-free survival included a hazard ratio of 0.27 in favor of LE as compared to PE treated tumors (95 percentiles= 0.099 to 0.709, P=0.02; Figure 2G).

**Figure 2:**
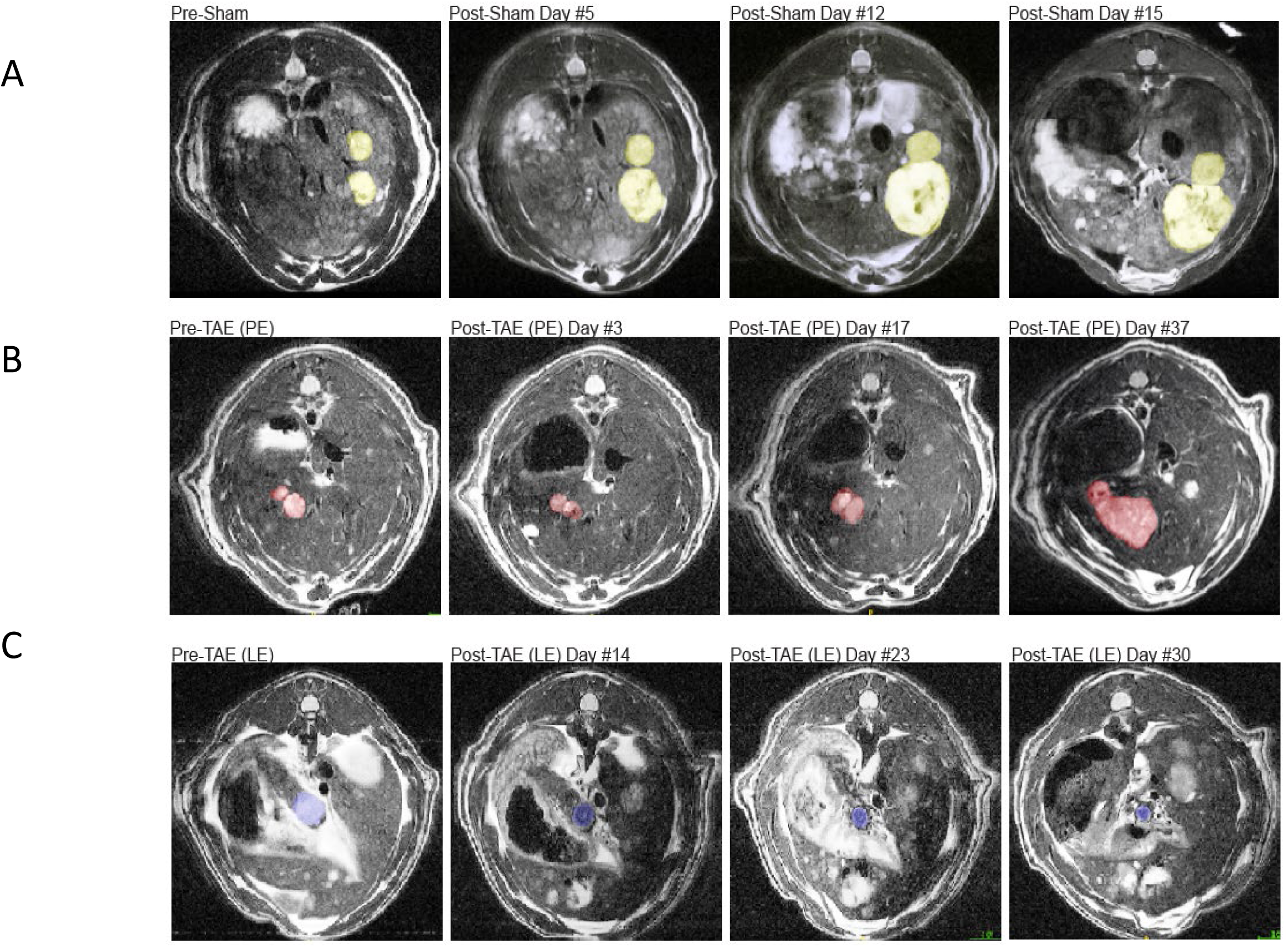
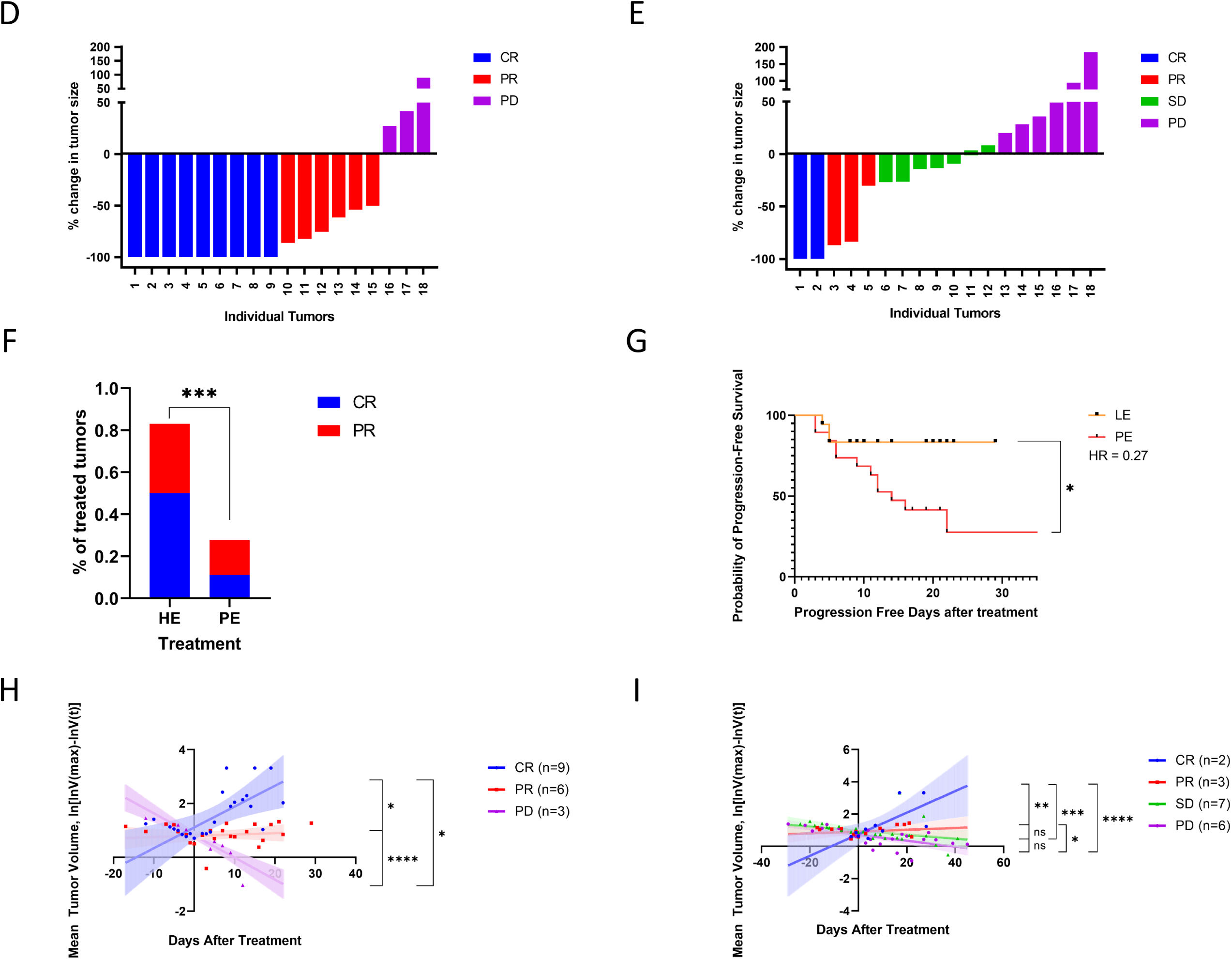
TAE with LE enables improved tumor targeting as compared to TAE with PE. Representative serial T2-weighted MR images target tumors prior to, and following, sham TAE (a, yellow), TAE with PE (b, red) and TAE with LE (c, blue). Bar graphs of RECIST responses for individual tumors treated with LE (d) and PE (e). (f) Bar graph of objective response rates following TAE with LE or PE. (g) Kaplan-Meier survival curve representing local progression-free survival for target tumors. Gompertzian transformed tumor growth curves stratified by RECIST response criteria for LE (h) and PE (i) treated tumors. CR = complete response. PR = partial response. SD = stable disease. PD = progressive disease. LE = liquid embolic. PE = particle embolic. HR = hazard ratio. * = P <.05, ** = P <.01, *** = P <.001, and **** = P <0.0001.

These findings were underscored by tumor growth kinetics following TAE. Linear regression of Gompertzian transformed tumor volumes provided specific growth rates for each cohort of tumors. LE TAE-treated tumors achieving a CR demonstrated a significantly reduced growth rate as compared to the PR or the PD groups (0.077 vs 0.005, P=0.02; and 0.077 vs -0.082, P=0.01, respectively Figure 2H;). LE TAE-treated tumors achieving a PR showed a significantly slower growth rate as compared to those demonstrating PD (0.005 vs -0.082, P<0.0001, Figure 2H). Similarly, PE TAE-treated tumors achieving a CR also demonstrated a significantly slower growth rate as compared to PR, SD, and PD groups (0.067 vs 0.006, P=0.0036; 0.067 vs -0.012, P=0.0007; 0.067 vs -0.018, P < 0.0001, respectively; Figure 2I). There were no significant differences in tumor growth rates for the PE TAE-treated tumors between the PR, SD, and PD groups (Figure 2I).

### Liquid Embolic Penetrates to the Level of Intratumoral Vasculature

Following selective TAE with LE, histological evaluation demonstrated deposition of polymerized liquid embolic within the tumor parenchyma and intratumoral vasculature (Figure 3A). Polymerized hydrogel was also identified in tubular and ovoid conformations within central and peripheral segments of tumor suggestive of vasculature; however, these areas did not stain positively for vascular markers including CD31 or SMA (Figure 3B). Polymerized LE was also discerned in necrotic areas of treated tumors (Figure 3C). Finally, diffuse intravascular and intraparenchymal punctae were identified in LE treated tumors consistent with inert, non-crosslinked PEG groups (21).

**Figure 3:**
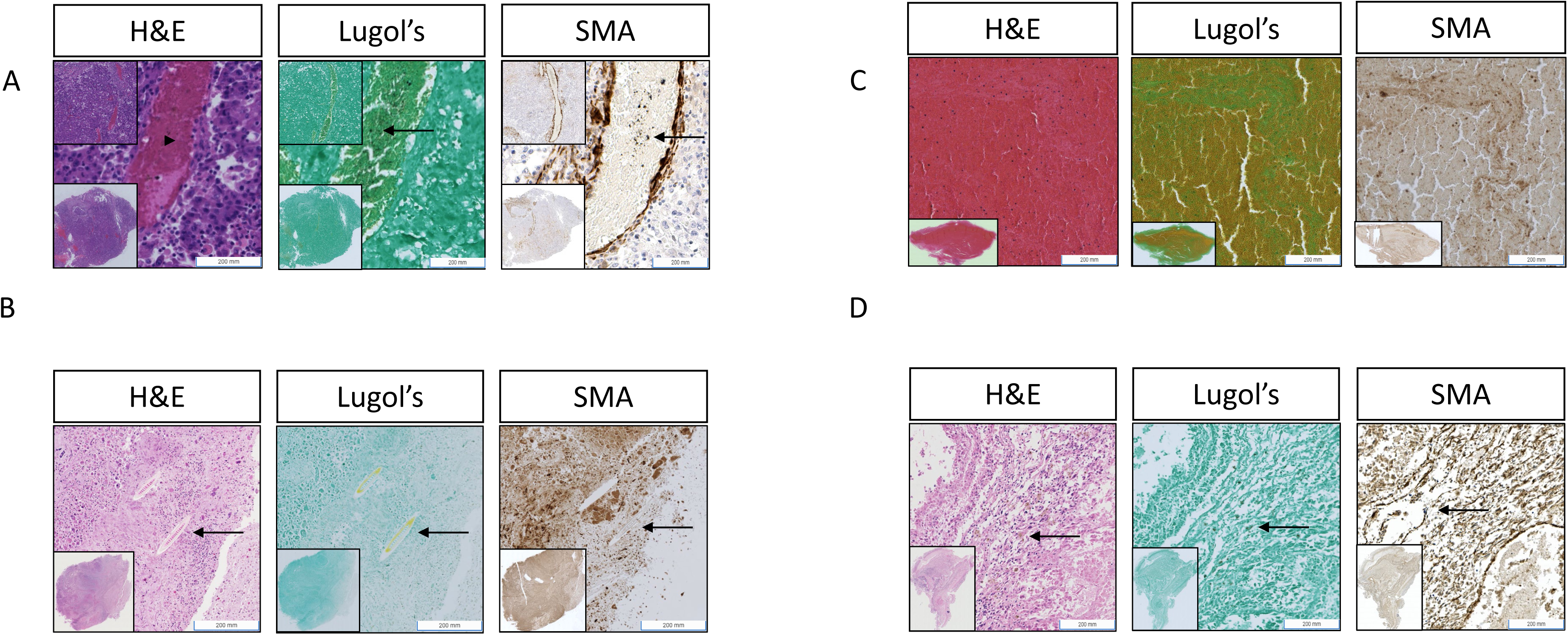
Histological evaluation of LE deposition within HCC and tumor parenchyma. (a) Polymerized LE staining yellow on Lugol’s stain deposited within the vascular bed with several black punctae of non-crosslinked PEG (black arrow). (b) Intratumoral polymerized LE in tubular configuration without surrounding structures staining positive for vascular markers (black arrow). (c) Lugol positive polymerized liquid embolic within necrotic tumor parenchyma (d) Intraparenchymal dark punctae consistent with non-crosslinked PEG particles (black arrow). SMA = smooth muscle actin. PEG = polyethylene glycol.

### TAE with Liquid Embolic Demonstrates Greater Tumor Necrosis at Early Time Points

To evaluate the mechanism underlying the observed differences in therapeutic response, H&E and pimonidazole staining were performed at 72 hours following treatment with LE, PE or SHAM therapy (Figure 4A). One-way ANOVA analysis demonstrated significant differences between the treatment groups with respect to % viable area, % necrotic area, and % necrotic and hypoxic area (P=0.0011, P=0.0010, and P=0.0004 respectively). Intragroup multiple comparison analysis showed that LE TAE resulted in a significantly lower % of viable tissue as compared to PE TAE (15.71±7.6 vs 43.10±.4, P=0.035), and significantly higher % necrotic tissue (84.48±7.7 vs 56.90±9.4, P=0.034) and combined % necrotic and hypoxic tissue (93.87±2.6 vs 63.39±9.5, P=0.019) (Figure 4B-E).

**Figure 4:**
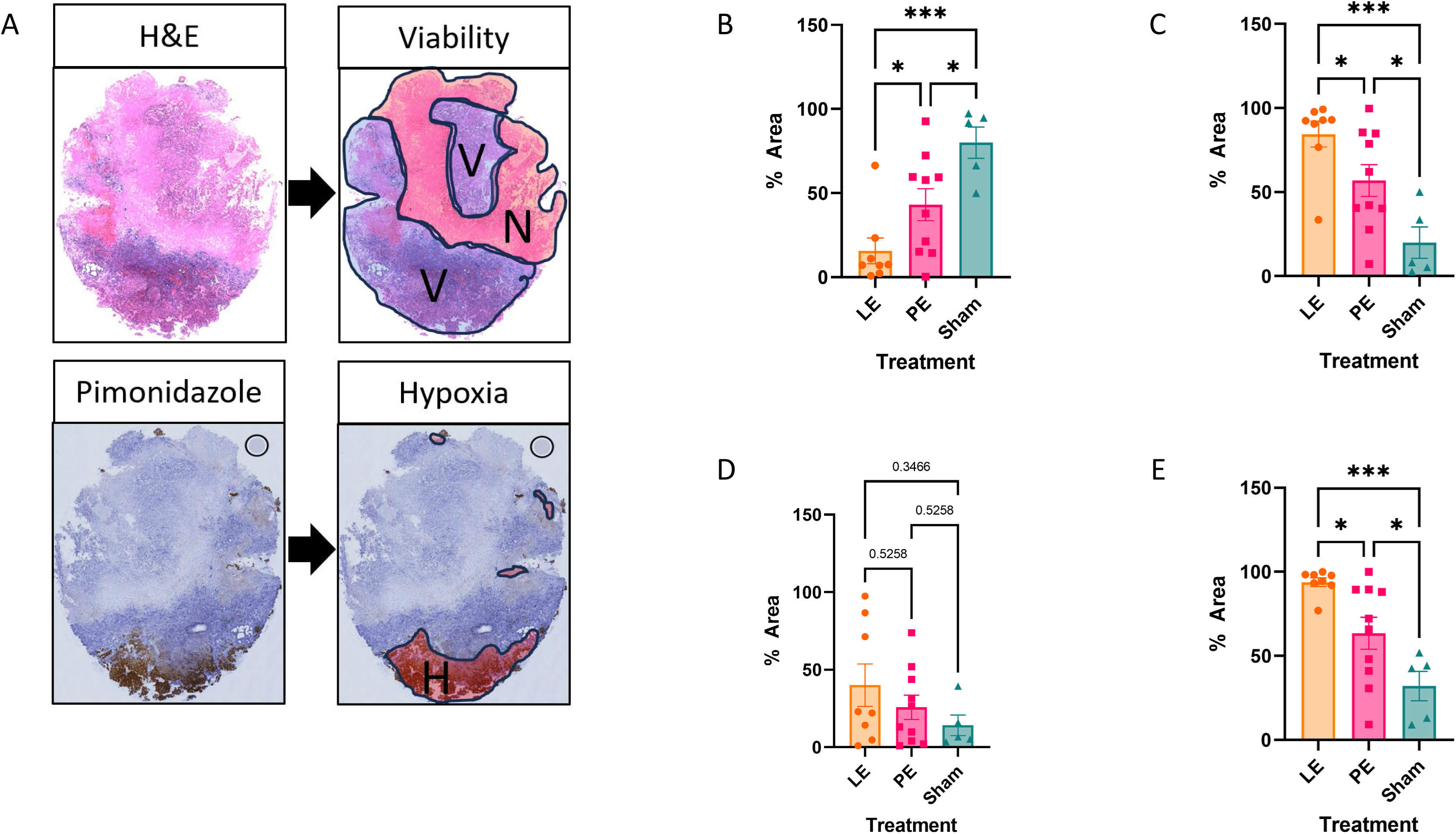
TAE with LE induces greater tissue necrosis as compared to TAE with PE or sham therapy. (a) Image J analysis identified areas of viability and necrosis across acquired H&E samples. Pimonidazole stained tumors were assessed for hypoxia within viable areas with (b) significant differences in the amount of viable tissue across treatment groups. (c) Bar graph demonstrating significant differences in the amount of necrotic tissue across treatment groups. (d) Bar graph showing a greater degree of intra-tumoral hypoxia in tumors treated with LE that did not reach statistical significance. (e) Bar graph demonstrating a significant difference in the amount of tissue that is both necrotic and hypoxic across treatment groups. LE = liquid embolic. PE = particle embolic. * = P <.05, *** = P <.001.

## Discussion

Leveraging an autochthonous HCC model in a background of hepatic cirrhosis that recapitulates vascular phenotypes observed in human disease, these data demonstrate differential antitumoral efficacy for a LE as compared to a PE including a markedly higher objective response rate (11). Consistent with these findings, TAE with LE enabled an improved local progression free survival, supporting the previously reported findings that LE may provide a more complete occlusion after formation of the intravascular casts(22). These findings were further supported by histological findings demonstrating extensive necrotic and hypoxic areas within LE-treated tumors indicative of more significant levels of tumor ischemia as compared to PE-treated tumors. Taken together, these findings emphasize a role for LE in optimizing ischemia-induced HCC cell death in the context of endovascular locoregional therapy.

TAE, often in combination with chemotherapy, remains a mainstay therapy for nonresectable HCC, yet the chemotherapeutic advantage remains unproven (23). Relevant prospective clinical trials and metanalyses point to non-superiority of transarterial chemoembolization (TACE) as compared to TAE in HCC patients, suggesting the importance of TACE-induced ischemia as a predominant mechanism of cytotoxicity over the effect of co-administered chemotherapeutics(24,25). The findings described herein underscore the importance of optimizing the induced ischemia to enhance tumor necrosis and hypoxia-induced cell death (26,27). In addition to enabling direct tumor targeting, ischemia has been shown to modulate the tumor microenvironment to induce targetable dependencies in surviving cancer cells as well as to modulate anti-tumor immunity (11,28–30). As such, robust and reproducible embolization techniques with durable effects hold the potential to also potentiate targeted therapeutics and facilitate immune priming for combination therapy (11,31).

Homogenously effecting ischemia is critically dependent on the ability of the applied embolic to occlude the intratumoral vasculature. The high rate of recurrence, underscored by data demonstrating limitations in the levels of hypoxia induced by TAE as well as vascular recanalization following treatment, emphasize a significant deficiency in the current embolic landscape, highlighting the need for improved embolics that can enable consistent and durable occlusion of the tumor vascular bed (2,5,6). Embolization of the proximal feeding vessels has been postulated to promote recanalization through small capsular feeding vessels that may be difficult to target using PEs (32). Furthermore, while particle-microvasculature mismatch has been shown to limit intratumoral penetration of existing PEs, matching embolic particles to the size of the target may not consistently address this issue as studies demonstrate conflicting results on the relationship between embolic size with anti-tumoral effectiveness (6,33,34). =.

LEs hold the potential to overcome these challenges through improved depth and completeness of embolization based on their unique vascular dynamics and ability to conform to target vessel geometry. The intravascular casts formed by LEs may be resistant to recanalization and less susceptible to the complexity or variability of intratumoral vascular anatomy (24,25). This potential was demonstrated in the current study given the significant improvement in necrotic and hypoxic tumor tissue following TAE with LE as compared to PE or sham therapy. The depth of penetration was demonstrated by the presence of LE within intra-tumoral vasculature as well as tumor tissue. In addition, the viscosity and polymerization time of newer LEs can be adjusted and tailored to unique applications, reducing potential non-target embolization and allowing optimal occlusion (22,35). In addition, LEs are not susceptible to the risk of intra-catheter deformity during delivery, or proximal clumping that can limit the efficacy of PEs (23,25). These features are underscored by the observed improvements in local progression-free survival and overall response rates for LEs as compared to PEs.

This study includes several limitations. Firstly, the observed disparity in response rates to LEs and PEs could be influenced by the PE to vessel size mismatch mentioned as well as the inherent heterogeneity that is characteristic of endovascular embolization and the associated influence of relatively small sample sizes. Secondly, the observed responses cannot be extrapolated to all LEs and PEs. Specifically, differences in the physical characteristics of PEs may influence efficacy. Finally, the location of the LE on histologic analyses was incompletely characterized as polymerized hydrogel in tubular confirmations could not be definitively determined to be intravascular. It is possible that these aggregates lie within the hepatic sinusoids which would not be expected to stain positive for arterial markers including SMA or CD31.

In conclusion, this study provides preclinical evidence supporting the potential of LEs to overcome limitations of PEs in inducing intra-tumoral ischemia in HCC. The LE utilized in this study affected a greater degree of tumor necrosis and hypoxia enabling superior tumor targeting as compared to the applied PE. These findings provide motivation for clinical studies in order to characterize their relevance to patients with HCC.

